# The endothelial scavenger receptor *stab2* is required for proper hematopoietic stem and progenitor cell development in the fetal blood stem cell niche

**DOI:** 10.64898/2026.07.07.737122

**Authors:** Gwendolyn M. Beacham, Zewde S. Ingram, Reema A. Elrefaie, Khaliun Enkhbayar, Zachary R. Zener, Leila Affini, Zainab N. Wasim, Maura C. Dodge, Sandeep Sreerama, Maria A. Serrano, Elliott J. Hagedorn

**Author notes:** Correspondence and material requests should be addressed to Elliott Hagedorn. These authors contributed equally (Gwendolyn M. Beacham, Zewde S. Ingram).

## Abstract

Hematopoietic stem and progenitor cell (HSPC) niches support lifelong production of blood and immune cells. Recently, we identified a gene signature unique to HSPC niche endothelial cells that is highly conserved across species and developmental time and includes the scavenger receptors *stab1/2* and *mrc1a*. Whether these receptors support HSPC development remains unclear. To investigate this, we used chemical inhibition and CRISPR mutagenesis in zebrafish and found that loss of *stab2*, and to a lesser degree *stab1*, reduced the number of embryonic *runx1*(+) HSPCs. Subsequent analyses of an additional HSPC marker (*cd41*) revealed an imbalance in the HSPC pool in *stab2* mutants, with reductions in *runx1*(+)*cd41*(+) and *runx1*(+)*cd41*(-) sub-populations containing stem cells and erythroid progenitors, respectively, the latter of which was most decreased. Our findings suggest stabilin scavenger receptors support HSPC development in the fetal niche, which could inform clinical strategies for culturing and expanding HSPCs.

## Introduction

Hematopoietic stem and progenitor cells (HSPCs) are formed during early embryonic development and sustain the lifelong production of blood and immune cells through their remarkable capacity to continuously self-renew while simultaneously generating differentiated hematopoietic lineages^1^. During a bone marrow transplant, this specific property of HSPCs is leveraged in what can be a curative treatment for patients with blood and immune disorders. In mammals, embryonic and adult HSPCs reside in highly specialized vascular niche microenvironments in the fetal liver and bone marrow, before and after birth, respectively. Different cell types within the niche are thought to support HSPC function, with sinusoidal endothelial cells (ECs) being a major contributor^2–5^. Although multiple adhesion and signaling molecules have been identified^6–9^, the full complement of EC-expressed niche factors that regulate HSPC development and activity remains incompletely understood.

The zebrafish has emerged as a robust vertebrate model for studying hematopoietic development where HSPCs and their supporting niche cells can be directly visualized with unparalleled resolution, in live, intact animals using a suite of fluorescent transgenic reporter lines^10,11^. Hematopoiesis and vascular biology are remarkably conserved between zebrafish and humans, sharing hematopoietic cell lineages as well as molecular pathways and cell-cell interactions that mediate the formation and maintenance of HSPCs during development. In zebrafish, the embryonic hematopoietic niche is housed in the ventral tail in a vascular plexus called the caudal hematopoietic tissue (CHT), which is thought to be the hematopoietic equivalent of the mammalian fetal liver^12^. Starting around 36 hours post-fertilization (hpf), nascent HSPCs coming from their site of birth in the aorta-gonad-mesonephros (AGM) region colonize the CHT where they reside and expand over the course of several days before migrating once again to the adult HSPC niche in the kidney marrow (hematopoietic equivalent to the mammalian bone marrow)^12,13^. Previously, we identified a 29-gene expression signature unique to the ECs of hematopoietic niches–both fetal and adult, in zebrafish and mice^14^. A predominant feature of this signature is the expression of several genes with functions related to endothelial scavenging. This includes the specific scavenger receptors *stabilin-1* (*stab1*)*, stabilin-2* (*stab2*) and *mrc1a*. Recent studies have shown that ECs in the CHT indeed exhibit a robust scavenger receptor-dependent endocytic activity that is involved in the clearance of macromolecules, lipid nanoparticles and cellular debris from circulation^15–18^. Although *stab1* and *stab2* have been previously implicated in regulating cellular interactions between hematopoietic and endothelial cells^19–23^, the potential functions of these receptors in the fetal HSPC niche with regards to HSPC development, have not been examined.

In this study, we systematically investigate the functional requirements of endothelial-expressed scavenger receptors in the HSPC niche of the zebrafish embryo. We show that conservation of these scavenger receptors’ expression in hematopoietic niche ECs extends to the human fetal liver, then use tools for chemical and genetic perturbation, paired with live cell imaging in zebrafish, to identify a specific requirement for *stab2* in promoting hematopoietic development. Although the cellular and structural organization of the CHT remain intact, we show that loss of *stab2* results in a specific reduction in HSPC sub-populations that include stem cells and erythroid progenitors, with the most severe decrease in the latter. Together, these findings have important implications for understanding the role of HSPC niche ECs in regulating hematopoietic development in the fetal niche.

## Results

### Conservation in the hematopoietic niche EC expression of scavenger receptors extends to the human fetal liver

In our recent study, we identified a conserved gene expression signature unique to the ECs of the fetal and adult hematopoietic niches of zebrafish and mice^14^. Amongst the most highly and selectively expressed genes were the scavenger receptors *stab1, stab2* and *mrc1a* (*Mrc1* in the mouse). Work by others have shown that *STAB1* and its paralog *STAB2* are expressed by sinusoidal ECs of the human bone marrow^2,24,25^. To determine whether these receptors are also expressed in the fetal hematopoietic niche in humans, we analyzed an existing publicly available scRNA-seq dataset of human fetal liver tissue^26^. We found that transcripts for *STAB1, STAB2* and *MRC1* were expressed in the *KDR*-positive endothelial cluster (**Figure 1A** and **Figure S1A**). Notably, of the three receptors, *STAB2* appeared to be the most highly and specifically expressed within the fetal liver ECs. *STAB1* and *MRC1*, by contrast, were also expressed broadly by an *MPEG1-*positive population that includes both monocytes/macrophages and Kupffer cells (**Figure 1A** and **Figure S1A**). Neither *STAB1*, *STAB2* nor *MRC1* were noticeably expressed by cells within the cluster that contained HSPCs (**Figure S1A**). Published data from independent scRNA-seq analyses of the human fetal liver showed similar expression^27,28^. These results confirm that the fetal hematopoietic niche EC expression of these specific scavenger receptors is indeed conserved in humans, with *STAB2* being particularly enriched.

**Figure 1.**
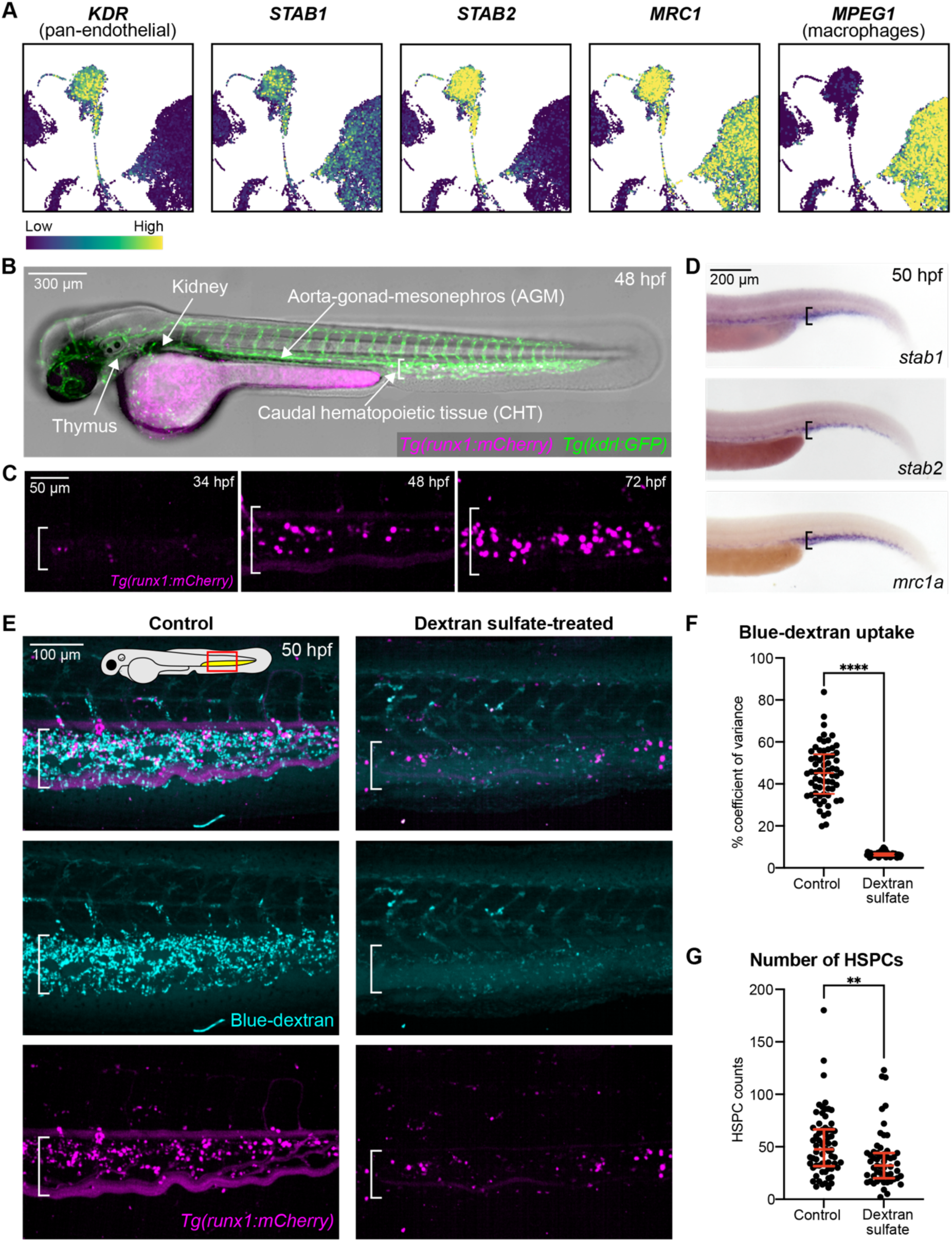
Endothelial-expressed scavenger receptors are required for proper numbers of *runx1*(+) HSPCs in the fetal hematopoietic niche. **A.** UMAPs of previously published single-cell RNA-sequencing data from human fetal livers (7–17 post-conception weeks) with a spectral scale indicating the normalized expression of the selected genes, adapted from Popescu et al., *Nature* 2019^26^. **B.** A zebrafish embryo expressing *runx1:mCherry* (HSPCs) and *kdrl:GFP* (all vascular ECs). Bracket denotes location of CHT in this and all subsequent figures. **C**. Images show *runx1:mCherry*(+) HSPC localization in the CHT (fetal HSPC niche) at three developmental stages. **D.** Whole-mount *in situ* hybridization (WISH) of *stab1*, *stab2* or *mrc1a* in *slc24a5* (control) crispants. **E.** Representative images of intravenously injected dextran-blue and *runx1:mCherry*(+) HSPCs in the CHT of control and dextran sulfate treated embryos. **F–G.** Quantification (percent coefficient of variance) of dextran-blue uptake (F) and *runx1:mCherry*(+) HSPC counts (G). For dot plots in this and all subsequent figures, each dot represents one zebrafish embryo. Data are from three pooled biological replicates in this and all subsequent figures, unless otherwise noted. Mann-Whitney test, error bars represent median ± interquartile range. **p<0.01, ****p<0.0001.

### Scavenger receptor inhibition reduces the number of *runx1*(*+*) HSPCs in the CHT

The potential role(s) of these endothelial-expressed scavenger receptors in supporting HSPCs in their niche, particularly *in vivo*, remains largely unexplored. Our previous studies examined hematopoietic niche EC gene expression in the CHT at 72 hpf^14^. As HSPC numbers increase significantly in this tissue between 36–72 hpf^11,13,29^ (**Figure 1B–C**), we used whole mount *in situ* hybridization to analyze the expression of *stab1*, *stab2* and *mrc1a* at an earlier stage of development. At 50 hpf, we observed strong expression of all three genes within the venous endothelial vasculature of the CHT (**Figure 1D**), consistent with what has been reported by other groups^17,30^. The developmental period during which HSPCs are housed in the fetal niche is thought to be critical for the expansion, maintenance/quality control and hematopoietic output of these specialized cells^12,31–33^. We therefore next sought to determine whether loss of function of these receptors might impact hematopoietic development. To test this, we utilized the *runx1+23:NLS-mCherry* transgenic zebrafish line (hereafter referred to as *runx1:mCherry*), which have nuclear-localized mCherry-labeled HSPCs that are capable of engraftment and expansion in the hematopoietic niches of donor animals following both adult-adult and embryo-embryo transplants^13^. We intravenously injected *runx1:mCherry* embryos with a broad scavenger receptor competitive inhibitor, dextran sulfate^34,35^. Dextran sulfate was intravenously injected into embryos via the duct of Cuvier at 36 hpf, a time point when nascent HSPCs from the AGM begin to colonize the CHT. To confirm scavenger receptor inhibition, we co-injected 10 kD cascade blue-conjugated dextran (dextran-blue) and fluorescein-conjugated hyaluronic acid (fluorescein-HA). These fluorescently conjugated macromolecules are known to be internalized by CHT ECs in a scavenger receptor-dependent manner^15,17^. To quantify endocytic uptake by CHT ECs we used spinning disk confocal microscopy and quantitative image analysis, measuring fluorescence within the CHT at 50 hpf. Consistent with previous studies^17^, we found that dextran sulfate treatment substantially reduced endocytic uptake of both dextran-blue and fluorescein-HA by the CHT ECs (**Figure 1E–F** and **Figure S1B–C**). In these same animals, we quantified the number of *runx1:mCherry*(+) HSPCs and observed a significant reduction in the dextran sulfate-treated animals compared to water-injected sibling controls (**Figure 1E&G**). Together, these results suggest that the activity of a fetal hematopoietic niche EC-expressed scavenger receptor(s) is required for normal HSPC development.

### HSPCs in the fetal niche are reduced in the absence of *stab2*

Since dextran sulfate is a broad inhibitor thought to target multiple scavenger receptors, we set out to evaluate each receptor individually. To this end, we adapted a previously published rapid F0 CRISPR whole-embryo knockout approach^36,37^. For each scavenger receptor gene, four top-ranked CRISPR guide sequences with predicted low off-target activity were selected based on the established criteria^36^ and used to synthesize single-guide RNAs (sgRNAs). Injection mixes containing all four ribonucleoprotein complexes (each of the four sgRNAs in-complex with Cas9 protein) were injected into individual zebrafish embryos at the one-cell stage to generate ‘crispants’ (F0 CRISPR knockouts). Using this platform, we generated individual crispants for *stab1*, *stab2* or *mrc1a.* As a negative control, we knocked out *slc24a5*, a gene whose loss generates the *golden* phenotype with a lack of pigmentation^38^. This strong phenotypic read-out also allowed us to confirm efficacy of our CRISPR/Cas9 reagents. Efficient targeting of the scavenger receptors was confirmed by reduced endocytic uptake of dextran-blue (*stab2* and *mrc1a*) and fluorescein-HA (*stab2*), whole mount *in situ* hybridization for transcripts of targeted genes (frequently reduced in crispants, likely due to nonsense-mediated decay) and/or next generation sequencing of target loci (**Figure S2A–F**). Imaging and quantification of *runx1:mCherry*(*+*) HSPC numbers at 50 hpf revealed that while *stab1* crispants had a modest reduction in HSPCs in the CHT compared to controls, *stab2* crispants had a large deficit in HSPCs (**Figure 2A–B**). Loss of *mrc1a*, by contrast, seemed to have no effect on HSPC numbers (**Figure 2A–B**). To further dissect the requirements of *stab1* and *stab2*, we knocked out both genes simultaneously, using half the amount of sgRNAs targeting each gene (to avoid toxicity) compared to our original CRISPR mixes. At these reduced doses, targeting *stab1* alone did not appear to have a significant effect on the number of HSPCs while targeting *stab2* alone still resulted in significantly fewer HSPCs in the CHT compared to controls (**Figure 2C–D**). The combined loss of *stab1* and *stab2* (*stab1/2* double crispants) resulted in HSPC numbers similar to in the *stab2* crispants (**Figure 2C–D**). We saw no decrease in viability after knocking out both genes, consistent with previous reports^15^. Of note, using half the amount of sgRNAs still appeared to disrupt gene function as uptake of dextran-blue by the CHT ECs was reduced in *stab2* and *stab1/2* double crispants (**Figure S2E–F**). We next longitudinally tracked *stab2* crispants over time and observed fewer *runx1:mCherry*(*+*) HSPCs in the CHT at 50, 60 and 72 hpf, and in the thymus (the site of embryonic T cell lymphopoiesis) and larval kidney marrow at 5 dpf compared to controls (**Figure 2E–G**). Supporting specificity of the CRISPR targeting, a similar decrease in HSPC numbers in the CHT was observed with an independent, distinct set of four sgRNAs targeting *stab2* (**Figure S2H–J**) and after crossing the *runx1:mCherry* transgene into a previously published stable *stab2^ibl2^* mutant^17^ (**Figure 2H**). We also found that the temporal expression of a *stab2 422 bp:GFP* enhancer transgene^14^ paralleled the developmental dynamics of HSPC localization, being first expressed in the CHT, then (starting at ∼4 dpf) in the developing kidney marrow, with *runx1:mCherry*(*+*) HSPCs directly interacting with GFP(+) ECs in both tissues (**Figure S3**). Consistent with our previous expression studies^14^ and with the human fetal liver scRNA-seq data^26^ analyzed in this study, we did not observe GFP expression in the HSPCs, only within the niche ECs (**Figures S1A** and **3**). Taken together, our results indicate a functional requirement for endothelial-expressed stabilin scavenger receptors, particularly Stab2, in regulating HSPCs in the fetal hematopoietic niche.

**Figure 2.**
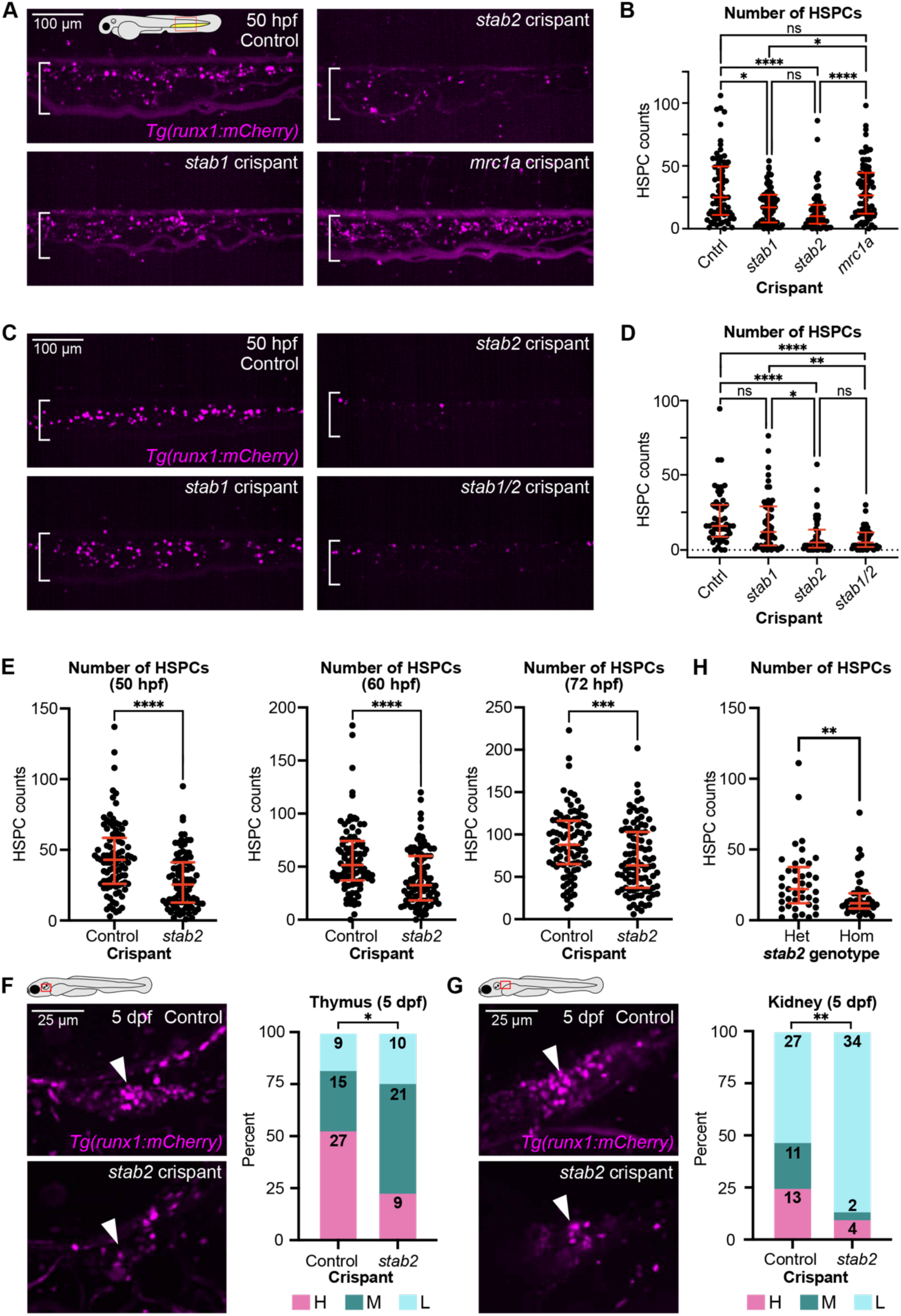
Loss of the specific stabilin scavenger receptor, *stab2*, reduces the number of *runx1*(+) HSPCs in the CHT. **A–D.** Representative images (A and C) and quantification (B and D) of *runx1:mCherry*(+) HSPCs in the CHT of the indicated crispants. **E.** Quantification of the number of *runx1:mCherry*(+) HSPCs as in B and D. Data shown at each developmental stage are from the same embryos over time. **F–G**. Images (left) and corresponding graphs (right) report quantification of *runx1:mCherry*(+) cells in the thymus (F) and kidney (G). High (H), medium (M) or low (L). The number of larvae for each category is indicated within the graphs. **H**. Quantification of *runx1:Cherry*(+) HSPCs in stable *stab2* homozygous mutants compared to heterozygous clutchmate siblings at 48 hpf. Control embryos for all crispant data are *golden* (*slc24a5*) crispants in this and all subsequent figures. For B and D, Kruskal-Wallis test, error bars represent median ± interquartile range. For E and H, Mann-Whitney test, error bars represent median ± interquartile range. For F and G, Chi square test. *p<0.05, **p<0.01, ***p<0.001, ****p<0.0001, n.s., not significant.

### Overall tissue structure and endothelial gene expression within the CHT are minimally affected by loss of *stab2*

A previous study in the zebrafish observed that *stab2* knockdown led to reduced Erk phosphorylation and altered arterial-venous differentiation, although this work was done using morpholinos, before CRISPR was widely available^39^. To confirm whether the developmental niche tissue was properly organized in our *stab2* crispants, we utilized high-resolution imaging of *kdrl:mCherry*(+) ECs (labels all blood vessels) in the CHT at 50 hpf. This analysis revealed that vessel architect was largely normal; while the CHT plexus in *stab2* crispants was slightly shorter in length compared to in controls, it was slightly taller in the dorsal-ventral axis, resulting in a total CHT niche area similar to controls (**Figure S4A–D**). Consistent with this, at 50 hpf, primitive macrophages and neutrophils were localized normally to the CHT in the absence of *stab2* (**Figure S4E–F**). Lastly, to assess whether loss of *stab2* altered other highly expressed niche endothelial genes, we examined an *mrc1a 125 bp:GFP* enhancer transgene^14^ and performed whole mount *in situ* hybridization for other CHT niche EC marker genes^14^ at 50 hpf. In each case, gene expression was unchanged or minimally affected in *stab2* crispants (**Figure S4G–H** and **Figure S2D**). Together, these results suggest that the reduction in HSPC numbers in the CHT of *stab2* mutants is not caused by significant morphological defects in the CHT tissue or changes to CHT niche EC identity.

### HSPC formation in the AGM and cellular dynamics of remaining HSPCs in the CHT appear normal in *stab2* mutants

To better understand what caused the reduction in HSPC numbers in the absence of *stab2*, we first examined the initial specification and formation of HSPCs in the AGM. Time-lapse imaging at 36 hpf revealed that the endothelial-to-hematopoietic budding of nascent *cd41:GFP; kdrl:mCherry* double positive HSPCs from the ventral dorsal aorta, was normal in *stab2* crispants (**Figure S4I**). To determine whether the deficit in HSPC numbers in the absence of *stab2* could be driven by reduced proliferation and/or increased cell death, we did a 12-hour (48–60 hpf) EdU labeling and TUNEL staining on fixed embryos at 50 hpf, respectively. In both assays, no significant differences were detected between control and *stab2* crispants at the developmental stages that were assessed (**Figure S4J–K**). Since Stab2 has previously been shown to facilitate adhesion between blood cells and ECs^20,24,40^, we tested whether loss of *stab2* in CHT niche ECs might alter HSPC migration by performing time-lapse imaging of *runx1:mCherry*(+) HSPCs as they arrived in the CHT. Despite being fewer in numbers, detailed cell tracking of individual HSPCs from ∼52–54 hpf showed no difference in niche residency time, suggesting HSPC migration dynamics over this 2 hour period were similar in *stab2* crispants compared to controls (**Figure S4L**). Altogether, these results suggest that the reduced HSPC numbers in *stab2* mutants are not driven by significant defects in HSPC formation in the AGM, alterations in proliferation/apoptosis or dysregulated migration.

### Loss of *stab2* reduces HSPCs labeled by *runx1+23* enhancer transgenes

In the zebrafish, multiple distinct fluorescent reporter transgenes are used to label HSPCs. To determine if the reduction in *runx1:mCherry*(*+*) HSPCs in *stab2* mutants was observed for HSPCs labeled by other commonly used markers, we generated *stab2* and control crispants in *cd41:GFP* and *cmyb:GFP* transgenic embryos. We imaged and quantified cell numbers using the same approach as for the *runx1:mCherry*(+) HSPCs. Surprisingly, we found that the number of *cd41:GFP*(*+*) and *cmyb:GFP*(*+*) HSPCs in 50 hpf embryos were similar between *stab2* crispants and controls (**Figure 3A–D**). Although there is overlapping signal between the different HSPC markers^13^, it is widely accepted that there is also additional transgene-specific labeling. We hypothesized that the *runx1+23* enhancer element might drive expression in a sub-population of HSPCs not marked by the other transgenes that is specifically affected by a loss of *stab2*. To test this, we first generated a new, independent *runx1:lifeact:mCherry* transgene using the same *runx1+23* mouse enhancer and beta-globin minimal promoter construct that was used to create the *runx1:mCherry* transgene (**Figure 3E**). The number of *runx1:lifeact-mCherry*(*+*) HSPCs was indeed reduced in *stab2* crispants compared to controls (**Figure 3F**), suggesting that these two *runx1* transgenes label a unique population of HSPCs that is particularly impacted by a loss of *stab2*.

**Figure 3.**
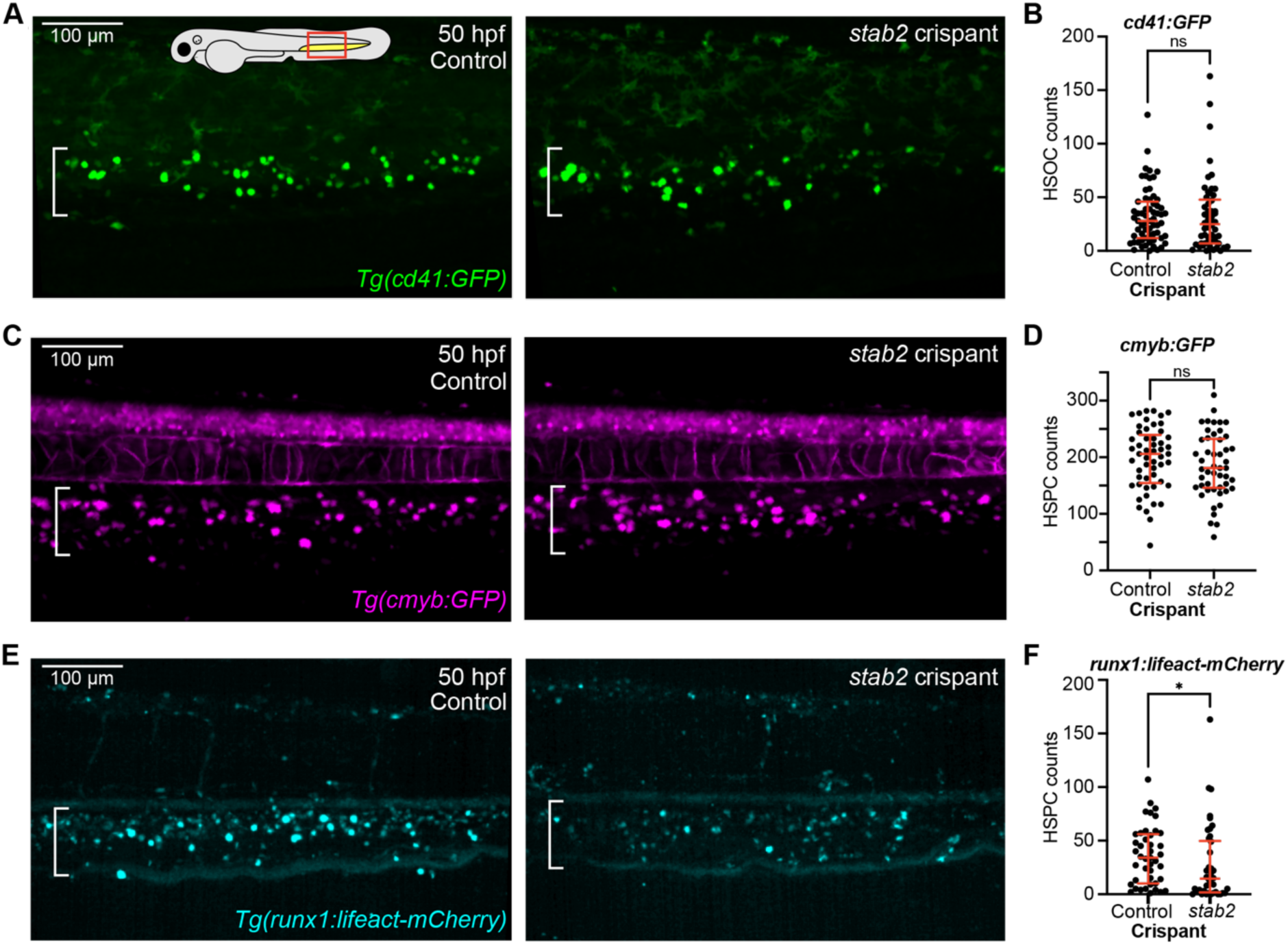
HSPCs labeled by *runx1+23* enhancer-driven transgenes, but not *cd41:GFP* or *cmyb:GFP*, are reduced in *stab2* mutants. Representative images (A, C, E) and corresponding quantification (B, D, F) report the number of HSPCs labeled by *cd41:GFP* (A and B, 48–50 hpf), *cmyb:GFP* (C and D, 50 hpf) and *runx1+23:lifeact-mCherry* (E and F, 50 hpf) in the CHT of *stab2* crispants or controls. Mann-Whitney test, error bars represent median ± interquartile range. *p<0.05, n.s., not significant.

### A *runx1*(*+*)*cd41*(*-*) erythroid progenitor population is dramatically reduced in the absence of *stab2*

To better resolve the specific HSPC population reduced in the absence of *stab2*, we crossed the *runx1:mCherry* line with *cd41:GFP*. Imaging of the CHT at 50 hpf revealed distinct populations of *runx1*(*+*)*cd41*(*+*) double positive and *runx1*(*-*)*cd41*(*+*) or *runx1*(*+*)*cd41*(*-*) single positive cells (**Figure 4A**), similar to what has been shown previously at 72 hpf^13^. To more precisely quantify the degree of overlap, we performed an object-based positional colocalization analysis, which showed that the double positive cells comprised 18.0% of the total cells labeled by either transgene, while the *cd41* and *runx1* single positive cells accounted for 54.1 and 27.9%, respectively (**Figure 4B**). To gain insight into the identity of the hematopoietic cells within these three populations, we used FACS to isolate live cells from double transgenic embryos expressing the *runx1:mCherry* and *cd41:GFP* transgenes (**Figure 4C**). To ensure sufficient cell numbers were collected, we sorted cells from embryos at 72 hpf. Sorted cells were then subjected to cytospin fixation, followed by May-Grünwald Giemsa staining for cytomorphological analysis. After imaging the slides, individual cells were classified into five distinct categories based on appearance (**Figure 4D**). This analysis revealed striking differences in morphology between the three populations: The majority of cells in the *cd41* single positive fraction were larger and had a distinctly vacuolated morphology, similar to what is often observed in cells of the myeloid lineage. A smaller percentage of the population appeared to be thrombocyte-like cells, consistent with previous reports that a portion of cells labeled by *cd41:GFP* are thrombocytes^41^. The double positive population contained both larger vacuolated cells and smaller distinctly round cells with a darker (blue) cytoplasm and low cytoplasm-to-cell size ratio. There was also a small amount of both thrombocyte- and erythroid-like cells. The *runx1* single positive population, by contrast, was very homogenous and comprised of cells with an erythroid progenitor-like morphology (**Figure 4D**), reminiscent of erythroid cells identified in previous studies of early hematopoietic development in the zebrafish embryo^42,43^.

**Figure 4.**
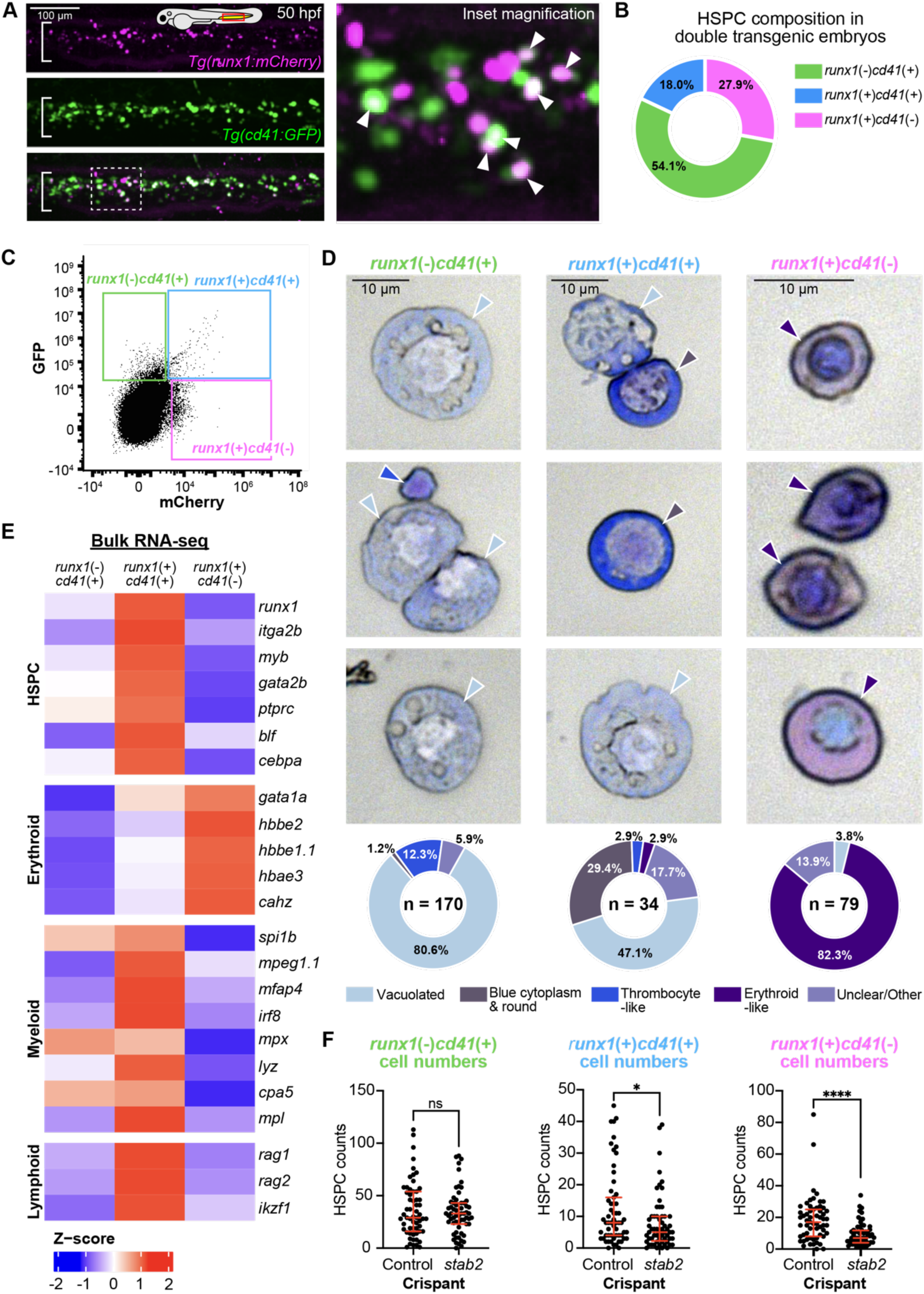
Hematopoietic sub-populations containing multipotent HSPCs and erythroid progenitors are reduced in the absence of *stab2*. **A.** Representative images from a single embryo (*slc24a5* crispant) expressing both *runx1:mCherry* and *cd41:GFP* transgenes. Dotted line indicates location of the inset magnification (right). White arrowheads point to HSPCs expressing both transgenes. **B.** Graph reports the percentage of HSPCs (3,995 cells from 60 control *slc25a5* crispants) that are labeled by *runx1:mCherry*, *cd41:GFP* or both transgenes in double transgenic animals. **C.** Representative flow cytometry plot and gating strategy for sorting HSPC sub-populations from 72 hpf *runx1:mCherry; cd41:GFP* (uninjected) double transgenic embryos. **D.** Representative images from two biological replicates of cytospin analyses of the sorted HSPC sub-populations, stained with May-Grünwald Giemsa and categorized based on morphology. Percentage of cells within each category is shown (bottom), with total number of cells scored (n) indicated. Colored arrowheads correspond to the designated category for each cell shown. **E.** Heat map showing *log_2_* transformed and Z-score normalized FPKM values of selected hematopoietic marker genes from one replicate of bulk RNA-seq in each of the three sorted sub-populations of HSPCs. Each row indicates the relative expression level of individual genes in each group compared to the group average. **F.** Number of HSPCs in each sub-population (identified by differential *runx1:mCherry* and *cd41:GFP* expression) in *stab2* and control crispants at 50 hpf. The control data in these experiments were used for the representative image in A and for the analysis shown in B. Mann-Whitney test, error bars represent median ± interquartile range. *p<0.05, **p<0.01, ****p<0.0001, n.s., not significant.

The expression of *runx1:mCherry* within erythroid cells is consistent with expression data reported by other groups^44,45^. Indeed, when we isolated RNA from the same three FACS-sorted populations for analysis by bulk RNA-seq, we identified a distinct expression profile for the *runx1* single positive population that was consistent with the cells being of an immature erythroid lineage (**Figure 4E** and **Table S3**). Although known red cell genes (e.g., *gata1a*^46^, embryonic globin genes [*hbbe2*, *hbbe1.1* and *hbae3*]^47,48^ and *cahz*^46^) were highly enriched, we also detected transcripts for genes known to be expressed by HSPCs, including *blf*^49^ and *cebpa*^50^ (**Table S3**)^50^, suggesting that cells in the *runx1* single positive population were recently derived from a progenitor and/or include immature erythrocytes. Consistent with this, in our cytospin analysis of the *runx1* single positive cells, we observed cells with varying degrees of nuclear compaction (**Figure 4D**), indicating the population likely contains cells at different stages of erythroid development^43,51^. While the double positive cells had some expression of these red cell genes, this erythroid signature was noticeably down in the *cd41* single positive cells compared to the other two populations (**Figure 4E** and **Table S3**). Within the double positive group, we observed strong expression for stem cell, myeloid and lymphoid genes (**Figure 4E** and **Table S3**), consistent with this population containing less differentiated HSPCs with multilineage potential (likely including true stem cells capable of self-renewal).

To determine whether *stab2* differentially regulates these distinct HSPC sub-populations (*runx1* or *cd41* single positive cells compared to double positive cells), we generated *stab2* crispants in the double transgenic embryos. When we compared the number of single and double positive cells in the CHT of *stab2* crispants and controls at 50 hpf, there was no difference in the number of *cd41* single positive cells (**Figure 4F**). By contrast, there was a reduction in the number of double positive cells in *stab2* crispants, and an even greater decrease in the number of *runx1* single positive cells (**Figure 4F**). Thus, the HSPCs that are specifically reduced in *stab2* mutants include both multipotent progenitors (double positive cells) and erythroid cells (*runx1* single positive cells), the latter of which are most reduced. All together, our findings indicate that stabilin scavenger receptors, most prominently *stab2*, support HSPC and erythroid cell development in the zebrafish hematopoietic niche.

## Discussion

Our data are consistent with a model in which stabilin scavenger receptors expressed by sinusoidal ECs in the fetal hematopoietic niche are required for HSPC and erythroid cell development. Stab1 and Stab2 are homologous transmembrane receptors that together comprise the family of fasciclin-like hyaluronan (HA) receptor homologs^52^. Both receptors are highly expressed by ECs of fetal and adult HSPC niches, in both zebrafish and mammals^14,19,25,53^, including in the human fetal liver, as we show in this study (**Figure 1A** and **Figure S1A**). In our zebrafish experiments using crispants, we found that loss of *stab1* resulted in a mild but statistically significant reduction in the number of *runx1:mCherry*(+) HSPCs in the CHT. While this reduction was not observed when we used half the amount of sgRNAs to knock out *stab1* (alone or in combination with a loss of *stab2*), our results suggest that *stab1* may also play a role in regulating HSPCs, albeit a more minor one than that of *stab2*. Interestingly, work by other groups has shown that Stab1 and Stab2 in the CHT endothelium exhibit partial redundancy in the clearance of anionic particles from circulation^15,17^. Consistent with this, we saw that uptake of 10 kD fluorescent dextran by CHT ECs is more reduced in *stab2* than in *stab1* crispants, but most reduced in embryos lacking both receptors (**Figure S2E**). By contrast, loss of the mannose scavenger receptor, *mrc1a*, which is also highly expressed by the ECs in HSPC niches, did not appear to impact HSPC numbers in the CHT (**Figure 2A–B)**. Our results therefore suggest that the stabilin scavenger receptors specifically (and most prominently *stab2*) are required for the balanced development of HSPCs in the hematopoietic niche of the zebrafish embryo.

Multiple distinct transgenes in the zebrafish embryo have been used as markers of HSPCs at various stages of hematopoietic development, from the posterior blood island and AGM through the CHT and kidney^10^. It is well-known within the field that different HSPC transgenes have overlapping expression patterns but also label distinct hematopoietic cell types. Even transgenes using the same regulatory element can label substantially different cell populations. Although the *runx1:lifeact-mCherry* line generated in this study appears to label a similar population of HSPCs compared to the *runx1:mCherry* transgene, the previously published *runx1:GFP* transgene (not used in this study but made using the same *runx1+23* enhancer) labels a much narrower population of HSPCs^13^, likely due to positional effects linked to the site of transgene integration, transgene silencing or a combination of the two. Indeed, despite there being fewer *runx1:mCherry*(+) HSPCs in the absence of *stab2*, we did not detect a significant difference in the total number of HSPCs labeled by the *cd41:GFP* or *cmyb:GFP* transgenes. We subsequently found that in *stab2* crispants, there was a mild reduction in the subset of *cd41:GFP(+)* cells that also expressed the *runx1:mCherry* transgene, while the *cd41:GFP*(+) *runx1:mCherry(-)* cells were largely unaffected. Indeed, our RNA-seq analysis revealed that these subsets of *cd41:GFP(+)* cells expressed different transcripts, indicative of distinct hematopoietic lineages. As the *cmyb:GFP* transgene is known to label hematopoietic cells more broadly than either *runx1:mCherry* or *cd41:GFP*, it is possible that the specific sub-population of HSPCs reduced in *stab2* mutants is labeled by *cmyb:GFP*, but that any reduction of these cells in *stab2* mutants is masked by the extensive labeling of other cells unchanged in the absence of *stab2*.

A subset of cells that highly express the *cd41:GFP* transgene (*cd41^high^*) have been characterized as thrombocytes, while the *cd41^low^* population is thought to contain more HSPC-like cells^41^. In our cytomorphological analysis, we observed thrombocyte-like cells in each of the *cd41*(+) populations but not in the *runx1* single positive population (**Figure 4D**), with the thrombocyte markers *mpl*^54^ and *itga2b*^41^ being most enriched in the double positive population (**Figure 4E**). We observed no correlation between brighter or dimmer *cd41:GFP*(*+*) cells overlapping with brighter or dimmer *runx1:mCherry*(+) cells (**Figure 4A**), at the developmental stages we analyzed (48–72 hpf). We found that the cells expressing both *cd41:GFP* and *runx1:mCherry* had the highest expression of stem cell/early progenitor marker genes (e.g., *runx1*, *myb* and *gata2b*) and this population was decreased in the absence of *stab2*. As both *cd41:GFP*(*+*) and *runx1:mCherry*(*+*) cells from the adult kidney marrow are independently capable of restoring the blood and immune system of irradiated recipient zebrafish^13,55^, it is not surprising that the sub-population of HSPCs labeled by both transgenes would include self-renewing stem cells with multilineage potential.

Hematopoiesis is thought to occur through multiple sequential but overlapping stages that are divided into ‘primitive’ and ‘definitive’ waves. Primitive hematopoiesis produces the initial erythrocytes, macrophages and neutrophils in the early embryo^56,57^. Soon after, during the first phase of definitive hematopoiesis, a transient population of erythromyeloid progenitors (EMPs), which are bi-potential and can differentiate into erythroid and myeloid but not lymphoid cells, emerge in a region of the ventral tail called the posterior blood island (PBI), which develops into the CHT. The second definitive and final wave of hematopoiesis commences with the emergence of AGM-derived HSPCs via an endothelial-to-hematopoietic transition from the ventral dorsal aorta starting at ∼36 hpf. These AGM-derived HSPCs migrate to the CHT where they proliferate and expand for several days before re-entering circulation and migrating to the thymus and kidney marrow (the sites of adult hematopoiesis). In zebrafish and mice, HSC-independent embryonic hematopoietic progenitors have been shown to sustain hematopoiesis longer than originally thought, potentially even into adulthood^45,58–60^. Whether the erythroid progenitors decreased in the absence of *stab2* are from HSC-independent or HSC-derived hematopoietic progenitors remains an open question. *Stab2* is indeed expressed by ECs in the PBI prior to colonization by AGM-derived HSPCs in the CHT (e.g. at 24 and 32 hpf^17,30^), and could potentially impact HSC-independent hematopoietic progenitors, like EMPs, at these earlier stages. Lineage tracing experiments in future studies, similar to ones used in recent studies by other groups^61,62^, could help to clarify the specific populations of HSPCs that give rise to these *runx1:mCherry*(*+*) erythroid progenitors that are most significantly reduced in the absence of *stab2*.

The most widely studied function of the endothelial-expressed stabilin scavenger receptors in tissues such as the spleen, liver, thymus and bone marrow, is to facilitate the internalization of a variety of ligands from circulation via clathrin-dependent endocytosis^63^. These ligands include, but are not limited to, apoptotic cells^64,65^, oxidized albumin^66^, phosphorothioate-modified antisense oligonucleotides (therapeutic agents)^67^ and modified low density lipoproteins^68^. Stab2 specifically also binds and internalizes hyaluronic acid^69^. Studies in the zebrafish embryo have investigated the scavenging function of these receptors in the CHT and found that Stab1 and Stab2 both readily mediate internalization of polyanionic nanoparticles, with Stab1 showing a preference toward the uptake of smaller particles compared to Stab2^15,17^. Whether this robust endocytosis driven by Stab1 and Stab2 relates to a function in balancing hematopoietic output in the fetal niche is a topic for further investigation. Interestingly, the conserved gene signature of HSPC niche ECs we previously identified includes several genes in addition to *stab1/2* and *mrc1a* that function in endocytosis and vesicle trafficking (e.g., *cltca*, *dab2*, *ap1b1* and *exoc3l2a*)^14^, suggesting these cellular pathways within the ECs might play a key role in supporting HSPCs. The stabilin scavenger receptors have also been shown to mediate the adhesion of migrating cells (e.g., lymphocytes and cancer cells) to the endothelium^20,70–72^, and an *in vitro* study identified a potential role for Stab2 in mediating the endothelial adhesion of HSPCs, specifically by binding to HA on the surface of HSPCs^19^. While our *in vivo* time-lapse imaging in the CHT of *stab2* crispants did not reveal a dramatic defect in adhesion/migration of *runx1:mCherry*(*+*) HSPCs (**Figure S4L**), the idea that Stab2 might indirectly or directly regulate HSPC-EC interactions or communications influencing HSPC development, remains an open hypothesis.

### Limitations of the study

The focus of this study was to determine whether endothelial-expressed scavenger receptors support HSPC development in the fetal hematopoietic niche. Whether or not the imbalance in the larval HSPC pool we observed in *stab1* and *stab2* mutants persists into adulthood will be important to clarify with future studies. Similarly, whether this imbalance in the HSPC pool has broader physiological consequences for the embryo or adult remains an open question. These stabilin scavenger receptors are highly conserved in mammals where they similarly are expressed by ECs of HSPC niches, including in the fetal liver. Future studies using human cells and mammalian models (e.g., mice) will be necessary, both for establishing the conservation of function and translating this work towards therapeutic contexts, which could include the *ex vivo* expansion of HSPCs prior to their transplantation into patients. Nonetheless, our study advances our fundamental understanding of endothelial expressed factors in the HSPC niche, demonstrating that stabilin scavenger receptors, especially *stab2*, are required for proper HSPC development during early embryogenesis.

## Methods

### Animals

Wild-type AB, *casper* or *casper-EKK* and the following transgenic and mutant zebrafish (*Danio rerio*) lines were used in this study: *runx1:mCherry [runx1+23:NLS-mCherry*]^13^, *kdrl(flk1):EGFP*^73^*, Stab2 422bp:EGFP*^14^*, stab2(ibl2)*^17^*, kdrl:mCherry [kdrl:Hsa.hras-mCherry]*^74^*, mrc1a 125bp:EGFP*^14^*, mpeg1:mCherry*^75^, *mpx:EGFP [zMPO:EGFP]*^76^*, cd41:EGFP*^41^, *cmyb:EGFP*^77^ and *runx1:lifeact-mCherry [runx1+23:lifeact-mCherry]* (this paper). Full transgene names are listed in brackets. All zebrafish were housed at Boston University School of Medicine and handled according to protocols approved by the Institutional Animal Care and Use Committee (IACUC) of Boston University. Adult zebrafish are cared for by the Laboratory Animal Science Center (LASC) of Boston University and maintained at 28.5°C with standard 14 h light, 10 h dark cycle on a recirculating system. Fish were bred by placing male-female pairs into a mating tank with dividers that were removed the following morning to allow natural spawning. Embryos were collected and grown in E3 media in an 28.5°C incubator. For all experiments, only healthy embryos with normal blood flow were used. To reduce plastic waste, plastic dishes were cleaned and reused using standard sterilization techniques whenever possible^78^.

### Analysis of human fetal liver scRNA-seq

Gene expression analysis of the human fetal liver was performed using a publicly available scRNAseq data set and online analysis software (accessible at https://developmental.cellatlas.io/studies/fetal-liver/dataset/192/cherita)^26^ using the Viridis color scale and the same visual scale adjustment across genes.

### Zebrafish intravenous injections

Embryos staged 36 and 48 hpf were immobilized with 1x tricaine (MS-222, pH 7.0, 160 mg/L) and placed on a 1.5% agarose pad with minimal E3 media. 1 nL of 0.01 mg/mL fluorescein-conjugated hyaluronic acid (fluorescein-HA; TdB labs) or 0.05 mg/mL cascade blue-conjugated dextran (10,000 MW) (blue-dextran; Invitrogen) was injected into the duct of Cuvier. Following injection, embryos were transferred to E3 without tricaine and incubated at 28.5°C until imaging. For dextran sulfate treatments, 1 nL of 10 mg/mL dextran sulfate (Sigma) or water control was co-injected with fluorescein-HA and blue-dextran into the duct of Cuvier of 36 hpf embryos.

### Whole-mount *in situ* hybridization (WISH)

Antisense *in situ* probes were generated by PCR amplification using cDNA, followed by reverse transcription with digoxygenin nucleotides (Roche) and T7 polymerase (Roche). **Table S1** lists all oligos used in probe generation. Zebrafish embryos (50 hpf) were euthanized by tricaine overdose, fixed in 4% paraformaldehyde (PFA; Santa Cruz Biotechnology) overnight at 4°C and dehydrated in methanol (stored at -20°C). Whole-mount *in situ* hybridization was performed as previously described^79^. Stained embryos were transferred to 80% glycerol for scoring and imaging, using a Nikon SMZ18 stereomicroscope equipped with a DS-Ri2 color camera controlled by the Extended Depth of Focus (EDF) module in NIS-Elements software (Nikon).

### CRISPR mutagenesis

All sgRNAs were designed using CHOPCHOP version 3.0.0 using the criteria described previously^36,37^. For each target gene, the top ranked four sgRNAs were selected and corresponding DNA oligos were purchased from Integrated DNA Technologies (IDT). **Table S1** lists all oligo sequences used for sgRNA synthesis. A fill-in PCR with a high-fidelity polymerase was performed to create the DNA templates for *in vitro* transcription that included the target gene specific crRNA and tracrRNA scaffold sequence. DNA templates were purified and transcribed using *in vitro* transcription (one per sgRNA) with T7 polymerase (MEGAscript T7 Transcription Kit). Each sgRNA was purified and the concentration and quality confirmed using nanodrop and gel electrophoresis. 500 pg of each sgRNA (2000 pg total) in-complex with 600 pg of Alt-R S.p. Cas9 Nuclease V3 protein (IDT) was injected into the cell of one-cell stage embryos. Injected embryos were incubated in E3 media at 28.5°C until analysis. In most F0 crispant experiments (as indicated in the figure legends), four sgRNAs targeting the pigmentation gene *slc24a5* were used for injecting sibling clutchmate controls. To knock out both *stab1* and *stab2* simultaneously, 250 pg of each of the four sgRNAs for each gene in-complex with 600 pg of Cas9 protein were injected. Clutchmate siblings lacking only *stab1* or *stab2* for comparison were generated by using 250 pg of each sgRNA targeting either *stab1* or *stab2* and 250 pg of each of the four sgRNAs targeting *slc24a5*. To confirm knockout efficiency in *stab2* crispants when a new batch of sgRNAs or Cas9 was used or when injected into a new strain, fluorescein-HA and blue-dextran were injected and uptake by CHT ECs was quantified.

### Genotyping

Embryos (2 dpf) were euthanized by tricaine overdose and genomic DNA was obtained via lysis in Taq polymerase buffer (made in-house) with 0.5 mg/mL Proteinase K (Invitrogen) at 65°C for 1 h with heat inactivation. Primers to genotype the *stab2^ibl2^* mutation are included in **Table S1**. The amplified DNA product was digested with *SmII* restriction enzyme (NEB) and analyzed using gel electrophoresis to distinguish wild-type (168 bp + 110 bp) and mutant (274 bp) alleles.

### Next Generation Sequencing (NGS)

Genomic DNA was extracted as described for Genotyping. PCR with Q5 polymerase (NEB) was performed to amplify a ∼250 bp long DNA fragment centered around the sgRNA (guide) cut site, for each of the four guides included in the F0 CRISPR knockout mix. Reverse primers included a unique 9 bp barcode to enable identification of individual embryos after NGS of pooled DNA amplicons. **Table S1** lists all oligo sequences used for NGS. Successful amplification was verified by gel electrophoresis. Amplicons to sequence the four guide cut sites in 2–3 embryos were pooled in equal proportions. The pooled amplicons were purified and quality/concentration was confirmed using nanodrop and gel electrophoresis. Four separate reactions were used to sequence the four guide sites in a total of 11–12 embryos, per biological replicate. NGS was performed using the CRISPR sequencing platform at the Massachusetts General Hospital Center for Computational & Integrative Biology Core. Barcodes were decoded and mutational frequency and outcome (wild-type or frame-shift/non-frame-shift inducing mutations) assigned using a previously published and publicly accessible bioinformatic pipeline^80,81^. Any target sites with poor sequencing coverage (less than 100 read counts) were excluded from the data.

### EdU and TUNEL staining

For EdU labeling, 1 nL of 2.5 mM EdU (Invitrogen) in DMSO (or DMSO alone for controls) was injected into the yolk of 48 hpf embryos as previously described^82^. Injected embryos were transferred to E3 medium and incubated at 28.5°C for 12 h, euthanized by tricaine overdose and fixed in 4% PFA (Santa Cruz Biotechnology) overnight at 4°C. Embryos were washed three times for 15 min in 0.2% Triton-X in PBS and permeabilized with 1% DMSO and 1% Triton-X in PBS. EdU staining was performed using the Click-iT EdU Cell Proliferation Kit for Imaging - Alexa 647 (Invitrogen) following the manufacturer’s instructions. Embryos were incubated in the dark at room temperature for 1 h then washed with 0.2% Triton-X in PBS, 5 times for 5 min each, and mounted for confocal imaging.

For TUNEL, 50 hpf embryos were fixed in 4% PFA overnight at 4°C, washed three times for 15 min each with 1% Triton-X in PBS and then permeabilized with proteinase K (0.01 mg/µL) for 15 min. Embryos were then refixed in 4% PFA for 10 min, washed with 1% Triton-X and blocked with 3% bovine serum albumin (BSA). Embryos were incubated with TUNEL stain mix (*In situ* Cell Death Detection Kit, Fluorescein; Roche) at 37°C in the dark for 1 hour before being washed twice and mounted for imaging. For a negative control, kit enzyme was not added, as per manufacturer’s instructions. For a positive control, embryos were treated with 0.5 mg/mL DNaseI for 20 minutes at room temperature prior to fixation.

### Microscopy

Zebrafish embryos were anesthetized with 1x tricaine and embedded in 0.8% low melting point agarose dissolved in E3 media on glass-bottom 6-well plates (MatTek). Imaging was performed using a Yokogawa CSU-W1 spinning disk mounted on an inverted Nikon Eclipse Ti2 microscope equipped with a full environmental enclosure maintained at 28.5°C. Within each experiment, the same imaging settings were used across all control and experimental groups and replicates.

All images of HSPCs, macrophages and neutrophils in the CHT were acquired using a 20X objective with a single field of view in the center of the CHT. For the whole-embryo image (**Figure 1B**) and imaging the CHT vasculature (*kdrl:mCherry*; **Figure S4A–D**), multiple fields of view were tiled together to increase the imaging area. Confocal z-stacks (150 µm) were acquired with a z-step size of 5 μm. For imaging the thymus and kidney, 250 µm z-stacks were used.

For timelapses of HSPC formation (*kdrl:mCherry; cd41:GFP* double positive embryos), 150 µm z-stacks collected every 15 min for 16 h were acquired using the 20X objective with a single field of view containing the AGM. For timelapses of *runx1:mCherry*(*+*) cell migration in the CHT, 60 µm z-stacks with 7 µm z-step were acquired every 30 sec for 2 h using the 20X objective with a single field of view in the CHT.

### Image analysis

Whenever possible, data was blinded prior to scoring. The spots function in Imaris (Oxford Instruments) was used to quantify HSPC numbers (*runx1:mCherry*, *cd41:GFP; cmyb:GFP and runx1:lifeact-mCherry*; spot size of 5.5 µm), macrophages (*mpeg1:mCherry*; spot size of 10 µm) and neutrophils (*mpx:GFP*; spot size of 10 µm). Spots were only identified and counted for each image within a user-defined region of interest that contained the CHT. Manual adjustment/correction of spots assignment was minimal. The same spots quality threshold was used for analyzing control and treatment groups. Due to variable expression of the *runx1:mCherry*, *cd41:GFP*, *mpeg1:mCherry* and *mpx:GFP*, embryos were not pre-screened prior to imaging and quantification. As such, a small number of transgene-negative embryos are expected to be present in some experiments but equally present in control and treatment groups. For *runx1:mCherry*, datasets were excluded from analysis when the median number of *mCherry*(+) cells in the CHT at 50 hpf was less than 15 for the control group as these clutches were deemed to have insufficient labeling of the HSPC compartment. For *cmyb:GFP*, embryos were pre-screened for transgene expression in the spinal cord at 48 hpf.

To quantify endocytic uptake of fluorescent ligands and *mrc1a 125bp:GFP* expression, maximum image projections were analyzed in Fiji/ImageJ^83^. A line with pixel width of 85 was drawn through the middle of the CHT of each embryo, using this same region of interest for each fluorescent channel. A background measurement outside the fish (line width of 85) was acquired to calculate the background-subtracted MFI measurement in the CHT. For the endocytosis assays, the mean fluorescence intensity (MFI) and standard deviation of the pixel intensity were quantified, and used to compute the Percent Coefficient of Variation (%CV; standard deviation/mean, multiplied by 100). %CV was used instead of over MFI alone due to high levels of ligands that remained in circulation in treatment groups.

For *runx1:mCherry*; *cd41:GFP* double transgenic animals, object-object statistics of defined spots (as described above) in Imaris were used to identify single and double positive cells. After a careful analysis of representative control images, *runx1:mCherry*(*+*) and *cd41:GFP*(*+*) spots were considered to be marking the same cell (double positive) if there was a minimum distance of 3.5 µm or less separating the spot centers.

To assess vascular morphology, measurements of *kdrl:mCherry* images were performed in Fiji/ImageJ. The length of the CHT was measured using a straight line from the start of the CHT vasculature (just posterior to the end of the yolk extension) to the point at which the artery transitions into a vein at the tip of the tail. The height of the CHT was measured in the widest section of the tail, from just below the artery to the bottom of the ventral most vein. The area of the CHT was measured using the freehand selection tool, and included the total area marked by *kdrl:mCherry* below the artery and posterior to the yolk extension.

Analysis of the AGM timelapses was performed using mp4 files generated in Imaris. The number of cells that were double positive for both transgenes (*kdrl:mCherry* and *cd41:GFP*) and budded from the endothelium were counted as nascent HSPCs.

For calculating the percent of *runx1:mCherry*(*+*) spots positive for EdU staining (imaging in fixed embryos), an object-object statistics analysis was performed as described above, using a minimum distance of 2 µm between spots for calling double positive spots. For TUNEL staining, prior to analysis, 3–5 *runx1:mCherry*(*+*) cells in the CHT (per embryo) were pre-selected blind for analysis. In Fiji/ImageJ, a region of interest (ROI) was drawn around the *runx1:mCherry*(*+*) signal using the freehand selection tool, and the TUNEL fluorescence intensity inside the ROI was measured and background-subtracted.

For HSPC migration timelapses (*runx1:mCherry*), the spots function in Imaris (same parameters as used for other analyses of this transgene) was used, paired with a tracking analysis (autoregressive motion). Only the tracks of spots that both entered and exited the field of view during the 2 hr timelapse were quantified. Tracks were edited to ensure that complete tracks for all spots that at any point during the timelapse met the quality threshold were included in the analysis. If a cell left the field of view or divided, the track was ended.

### Tol2 Transgenesis

To construct the *Tg(runx1:lifeact-mCherry)* line, an existing Tol2 vector containing *runx1+23*-driven mCherry was amplified using oligos that inserted the sequence for lifeact when recombined in a one-piece Gibson cloning reaction. **Table S1** lists all oligos and **Table S2** lists all plasmids used in the cloning. Embryos were injected with 25 pg plasmid DNA and 25 pg Tol2 mRNA at the one cell-stage as previously described^84^. To confirm successful integration, embryos were screened at 72 hpf for construct expression. Founders were grown to adulthood and outcrossed, and subsequent progeny were screened for stable integration of the construct.

### FACS sorting of *runx1:mCherry*(*+*) and *cd41:GFP*(*+*) cells

Double transgenic embryos (*runx1:mCherry*; *cd41:GFP*) were screened for expression of both transgenes. Embryos at 72 hpf were euthanized via tricaine overdose and prepared for FACS as previously described^14^. Briefly, embryos were minced using a razor blade, recovered in cold PBS and dissociated with liberase at 37°C for ∼40 min before filtering through a 40 μm cell strainer, pelleted by centrifugation and resuspended in FACS buffer (2% FBS; 1x PBS). DAPI (4′,6-diamidino-2-phenylindole; NucBlue Fixed Cell Stain ReadyProbes, Molecular Probes) was added to exclude dead cells from analysis. Sorting was performed on a BD FACSDiscover S8 spectral cell sorter with imaging capabilities. Gating was determined based on non-fluorescent and single color controls in aged matched embryos. Single and double positive cells were sorted into FACS buffer.

### Bulk RNA sequencing

Following sorting, cells were pelleted by centrifugation and total RNA was extracted using TRIzol (Invitrogen). Chloroform (15%) was added followed by vigorous shaking and centrifugation at 12,000 x g for 15 min at 4°C. GlycoBlue coprecipitant (Invitrogen) was added to the aqueous fraction and the sample was centrifuged at 12,000 x g for 20 min at 4°C. The RNA pellet was washed twice with 75% ethanol and resuspended in nuclease-free water. RNA concentration was quantified on Qubit 2.0 and samples were submitted for ultra-low input bulk mRNA sequencing (Novogene).

To generate a heatmap of differential gene expression, a subset of the expression matrix containing the FPKM values for selected genes of interest was *log_2_* transformed (*log_2_* (FPKM + 1)) in RStudio. After removing zero-variance genes, row-wise Z-score scaling was applied to each gene of interest to evaluate relative expression changes across samples. The resulting standardized matrix was visualized using the ComplexHeatmap R package.

### Cytospin analysis

Cytomorphological analysis of 5,000 sorted cells was performed using a Shandon Cytospin 3 centrifuge. Slides were stained using May-Grünwald Giemsa stain (Azer Scientific) according to the manufacturer’s instructions and imaged using a 100X oil objective on a Nikon Eclipse Ti2 inverted microscope equipped with a DsRi2 color camera. Images were scored and cells were categorized based on morphology. Any cells that could not clearly be placed in a single category were considered as ‘Unclear/Other.’ Cells that were too clumped together to clearly distinguish individual cell boundaries and objects that were not easily identifiable as cells were excluded from the analysis.

### Quantification and statistical analysis

All data shown are from three pooled biological replicates unless otherwise noted in the figure legend. GraphPad Prism software was used for all statistical analyses, with the statistical tests utilized noted in figure legends.

#### Acknowledgements

This work was supported by a Career Investment Award from the Department of Medicine at the Boston University Chobanian & Avedisian School of Medicine and Boston Medical Center (BU-BMC), an Early Career Scientific Research Award from the Association for the Advancement of Blood & Biotherapies, a Cancer Research Innovator Award from the BU-BMC Cancer Center, a BMC Bridge Funding Award, and NIH grants K01 DK111790 and R01 HL178832 to E.J.H. G.M.B. was a Merck Fellow of the Jane Coffin Childs Memorial Fund and Z.S.I. was supported by a graduate fellowship from the Carter Fund for Cancer Disparities at BMC and NIH grant F31 HL184822. G.M.B. and Z.S.I. were also supported by CTSI NIH/TL1 grant 5TL1TR001410-08. R.A.E. was supported by a Diversity in Cancer Research Award from the American Cancer Society and Z.R.Z. and Z.N.W. received support from the Boston University Undergraduate Research Opportunities Program. We gratefully acknowledge D. Ragoonanan, S. Song, Z. Amer and H. Wang for research and technical support, B. Neptune for operations support and A. Donnelly for grant support. We thank the labs of S. Cui, C. Heaphy, L. Lowery, R. Dries and G. Murphy for sharing space, equipment and reagents, and S. Avagyan for her generous assistance with the NGS analysis of crispants and for her helpful comments on the manuscript. Thank you to A. Chang and K. Parks for assistance scoring data and to S. Schulte-Merker for sharing the *stab2^ibl2^* mutant. Thank you to the aquatics staff at BU, in particular D. Walsh and H. Mullin, and to S. Brezinsky in the BU Flow Cytometry Core. The BD FACSDiscover S8 used in this study was purchased with NIH Shared Instrumentation grant S10 OD038346.

## Author contributions

G.M.B. and Z.S.I. designed the study, funded the project, performed experiments, interpreted the data and wrote and edited the manuscript. R.A.E., K.E., Z.R.Z., L.A., Z.N.W., M.C.D. and S.S. performed experiments and provided technical support. M.A.S. provided technical and conceptual support. E.J.H. designed the study, funded and supervised the project, interpreted the data and wrote and edited the manuscript. All authors reviewed the manuscript.

## Accession numbers

The previously published human fetal liver genomic data reported in this paper is deposited at ArrayExpress (accession code <u>E-MTAB-7407)</u>^26^. The zebrafish genomic data reported in this paper is being submitted to the NCBI Gene Expression Omnibus and the accession number will be forwarded upon receipt.

## Declaration of interests

The authors declare no competing interests.

## Notes

### Competing Interest Statement

The authors have declared no competing interest.

### Summary of Updates

We modified the order of data presented in Figure 4, and corresponding text in the Results section, and added additional data to Figure S1. We also incorporated additional literature into the Discussion.

