## Supplementary Material for "The endothelial scavenger receptor *stab2* is required for proper hematopoietic stem and progenitor cell development in the fetal blood stem cell niche"

### Supplemental information - Document S1

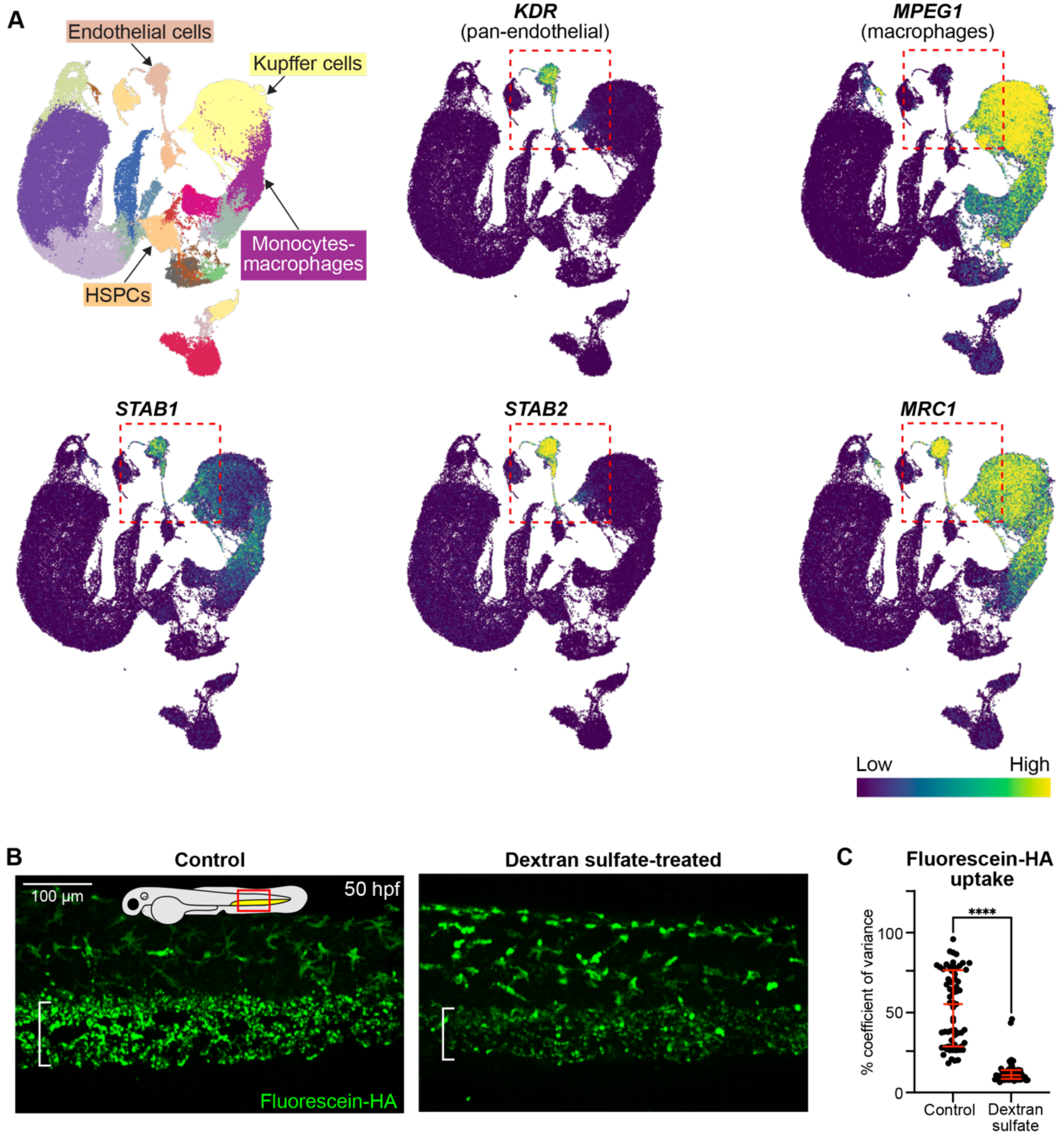


**Figure S1**

**Figure S1. Uncropped UMAPs for human fetal liver scRNA-seq data and quantification of HA uptake in the zebrafish CHT. A.** UMAP of human fetal liver scRNA-sequencing from Popescu et al., *Nature* 2019[26] with colors indicating cell type clusters (top left). Other UMAPs are uncropped versions of those shown in Figure 1A, with a red dotted line indicating the cropped region. **B–C.** Representative images (B) and quantification (C) of fluorescein-HA uptake in the CHT of the same embryos shown in Figure 1E. Bracket denotes location of CHT in this and all subsequent figures. For dot plots in this and all subsequent figures, each dot represents one zebrafish embryo, unless otherwise noted. Mann-Whitney test, error bars represent median ± interquartile range. ****$p<0.0001$.

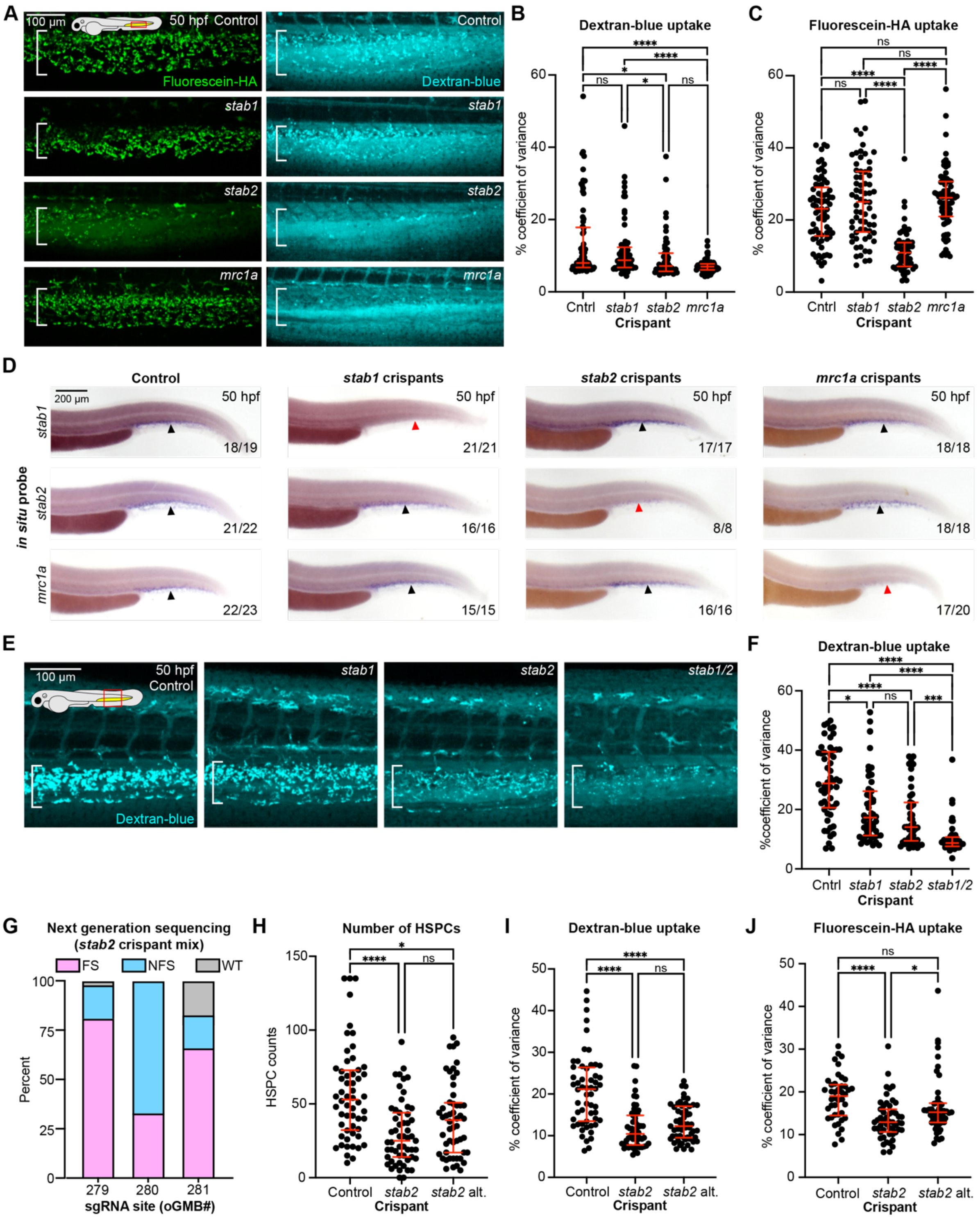
A
100 µm
50 hpf Control
Fluorescein-HA
Control
Dextran-blue
stab1
stab2
mrc1a
B
Dextran-blue uptake
% coefficient of variance
Cntrl stab1 stab2 mrc1a
Crispant
C
Fluorescein-HA uptake
% coefficient of variance
Cntrl stab1 stab2 mrc1a
Crispant
D
Control
stab1 crispants
stab2 crispants
mrc1a crispants
200 µm
50 hpf
in situ probe
stab1
stab2
mrc1a
18/19
21/21
17/17
18/18
21/22
16/16
8/8
18/18
22/23
15/15
16/16
17/20
E
100 µm
50 hpf Control
Dextran-blue
stab1
stab2
stab1/2
F
Dextran-blue uptake
%coefficient of variance
Cntrl stab1 stab2 stab1/2
Crispant
G
Next generation sequencing (stab2 crispant mix)
FS
NFS
WT
Percent
279 280 281
sgRNA site (oGMB#)
H
Number of HSPCs
HSPC counts
Control stab2 stab2 alt.
Crispant
I
Dextran-blue uptake
% coefficient of variance
Control stab2 stab2 alt.
Crispant
J
Fluorescein-HA uptake
% coefficient of variance
Control stab2 stab2 alt.
Crispant

**Figure S2**

**Figure S2. Additional characterization of *stab1*, *stab2* and *mrc1a* crispants. A–C.** Representative images (A) and quantification of dextran-blue (B) and fluorescein-HA (C) uptake in the CHT of control, *stab1*, *stab2* and *mrc1a* crispants (same embryos as in Figure 2A–B). **D.** Whole-mount *in situ* hybridization (WISH) of *stab1*, *stab2* or *mrc1a* in the indicated crispant background. Black arrowheads indicate staining in the CHT. Red arrowheads indicate reduced staining compared to in controls. The fraction of embryos that exhibit the shown phenotype across one biological replicate is indicated. **E–F.** Representative images (E) and quantification (F) of dextran-blue uptake in the CHT of control, *stab1*, *stab2* and *stab1/2* (double) crispants (same embryos as in Figure 2C–D). **G.** Percentage of frameshift causing (FS) and non-frameshift causing (NFS) CRISPR edits at three out of the four sgRNA cut sites in *stab2* crispants, identified using next generation sequencing. Sequencing across the 4th sgRNA site (not shown) produced low read counts (<100 per embryo) in the majority of embryos. Data are of two pooled biological replicates. **H–J.** Number of *runx1:mCherry* HSPCs (H) and quantification of dextran-blue (I) and fluorescein-HA (J) uptake in control and *stab2* crispants, compared to crispants generated with four different sgRNAs in *stab2* (*stab2* alt), at 50 hpf. The same embryos were quantified in H, I and J. Control embryos for all crispant data are *golden* (*slc24a5*) crispants in this and all subsequent figures, unless otherwise noted. Kruskal-Wallis test, error bars represent median ± interquartile range. *p<0.05, ***p<0.001, ****p<0.0001, n.s., not significant.

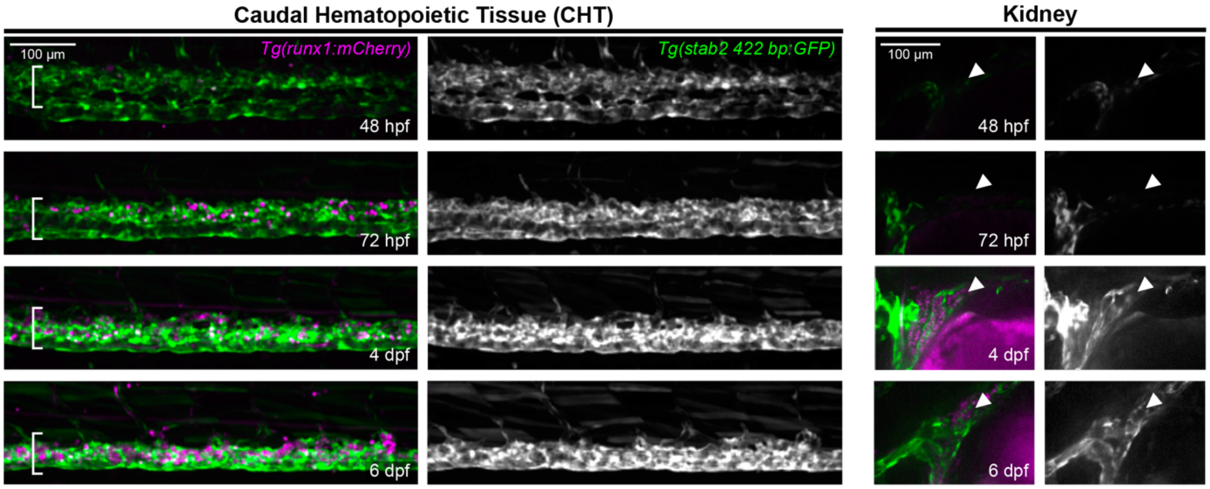


Figure S3

**Figure S3. A *stab2* enhancer transgene is expressed by ECs of the embryonic and developing adult hematopoietic niches.**

Representative images of *stab2* enhancer GFP and *runx1:mCherry* transgenes in the CHT (left) and larval kidney (right). GFP is shown in grayscale. White arrowheads indicate expression of the *stab2* enhancer GFP transgene in the kidney.

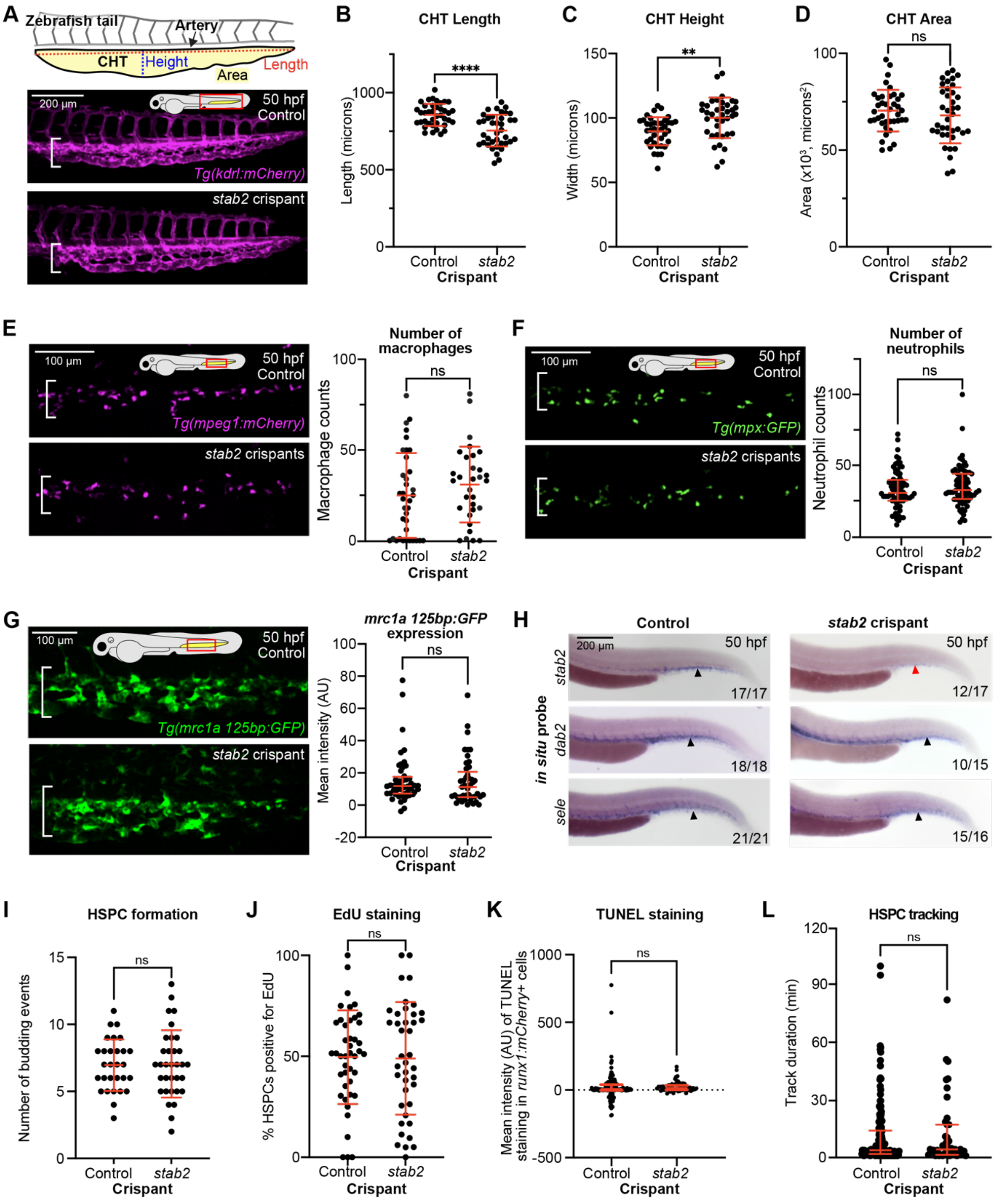
A
Zebrafish tail
Artery
CHT
Height
Area
Length
200 µm
50 hpf
Control
Tg(kdrl:mCherry)
stab2 crispant
B
CHT Length
****
Length (microns)
Control
stab2
Crispant
C
CHT Height
**
Width (microns)
D
CHT Area
ns
Area (x10^3, microns^2)
E
100 µm
Tg(mpeg1:mCherry)
stab2 crispants
Number of macrophages
Macrophage counts
F
Tg(mpx:GFP)
Number of neutrophils
Neutrophil counts
G
Tg(mrc1a 125bp:GFP)
mrc1a 125bp:GFP expression
Mean intensity (AU)
H
in situ probe
stab2
dab2
sele
17/17
12/17
18/18
10/15
21/21
15/16
I
HSPC formation
Number of budding events
J
EdU staining
% HSPCs positive for EdU
K
TUNEL staining
Mean intensity (AU) of TUNEL staining in runx1:mCherry+ cells
L
HSPC tracking
Track duration (min)

**Figure S4**

**Figure S4. Tissue structure, cellular composition and endothelial gene expression within the CHT remain intact in *stab2* crispants. A.** Schematic indicating CHT length, height and area measurements in the tail of the zebrafish embryo with representative images below. **B–D.** Graphs report the length (B), height (C) and area (D) of the CHT vascular plexus in 50 hpf crispants as depicted in A. **E–F.** Representative images (left) and quantification (right) of *mpeg1:mCherry*(+) macrophage (E) and *mpx:GFP*(+) neutrophil (F) numbers in the CHT of *stab2* crispants and controls. **G.** Representative images (left) and background-subtracted mean intensity (plotted, right) of a CHT EC-specific transgene, *mrc1a 125bp:GFP* in *stab2* and control crispants. **H.** WISH of niche EC genes selectively expressed in the vasculature of the CHT. Black arrowheads indicate staining in the CHT. Red arrowhead indicates reduced staining compared to in the control. The fraction of embryos that exhibit the shown phenotype across two pooled biological replicates is indicated. **I.** Number of nascent *cd41:GFP*(+) HSPCs that budded from the aorta-gonad-mesonephros region from 36–42 hpf in control embryos (uninjected) and *stab2* crispants. **J.** Percentage of *runx1:mCherry*(+) HSPCs positive for EdU staining in fixed embryos at 60 hpf (exposed to EdU from 48–60 hpf). **K**. Background-subtracted mean intensity of TUNEL staining in *runx1:mCherry*(+) HSPCs in 50 hpf fixed embryos (two pooled biological replicates). **L.** Track durations of *runx1:mCherry*(+) HSPCs during a 2 hr timelapse from ~52–54 hpf. In the dot plots in K and L, each dot represents one *runx1:mCherry*(+) HSPC (number of embryos for L: n = 9 for controls, n = 7 for *stab2* crispants; number of embryos for K: n = 25 for controls, n = 11 for *stab2* crispants). For B–D, E, I and J, Unpaired student's t-test, error bars represent mean ± standard deviation. For F, G, K, and L, Mann-Whitney test, error bars represent median ± interquartile range. **p<0.01, ****p<0.0001, n.s., not significant.

**Table S1| Olignonucleotides**

| Unique Identifier | Description | Category | Sequence | Source |
|---|---|---|---|---|
| oGMB1 | Universal primer for CRISPR sgRNAs | sgRNA template | AAAAGCACCGACTCGGTGCCACTTTTTCAAGTTGATAACGGACT AGCCTTATTTTAACTTGCTATTTCTAGCTCTAAAAC | This paper |
| oGMB279 | *stab2* guide 1 | sgRNA template | taatacgactcactataGGGGCAGATTCTAGTCAATGgttttagagctagaa | This paper |
| oGMB280 | *stab2* guide 2 | sgRNA template | taatacgactcactataGGCAACGGCATATGTAAAGAgttttagagctagaa | This paper |
| oGMB281 | *stab2* guide 3 | sgRNA template | taatacgactcactataGGAACTGGCAGGTGGCCTATgttttagagctagaa | This paper |
| oGMB282 | *stab2* guide 4 | sgRNA template | taatacgactcactataGGAGGTTGCTCAGCCCATGgttttagagctagaa | This paper |
| oGMB502 | *stab1* guide 1 | sgRNA template | taatacgactcactataGGGAGGTCCAAAGCAACGGCgttttagagctagaa | This paper |
| oGMB503 | *stab1* guide 2 | sgRNA template | taatacgactcactataGGCGATGGACTCTCGTGTTAgttttagagctagaa | This paper |
| oZSI17 | *stab1* guide 3 | sgRNA template | taatacgactcactataGGCGATGGCAGACGTTGCTAgttttagagctagaa | This paper |
| oZSI19 | *stab1* guide 4 | sgRNA template | taatacgactcactataGGTCGCCAGGTAGAGCTGTCgttttagagctagaa | This paper |
| oZSI24 | *mrc1a* guide 1 | sgRNA template | taatacgactcactataGGTCACCCCTCTCTGGAACCgttttagagctagaa | This paper |
| oZSI25 | *mrc1a* guide 2 | sgRNA template | taatacgactcactataGGGCTCAAAACATCAAGGACgttttagagctagaa | This paper |
| oZSI26 | *mrc1a* guide 3 | sgRNA template | taatacgactcactataGGAGGATTGTGTCCTCATCAgttttagagctagaa | This paper |
| oZSI27 | *mrc1a* guide 4 | sgRNA template | taatacgactcactataGGAGTCGCTGGCAGGTGTATgttttagagctagaa | This paper |
| oGMB538 | *stab2* guide 1, alt | sgRNA template | taatacgactcactataGGGTGTCAAGTTCATAAATAgttttagagctagaa | This paper |
| oGMB539 | *stab2* guide 2, alt | sgRNA template | taatacgactcactataGGGAACTGAGATCCTCTGGGgttttagagctagaa | This paper |
| oGMB540 | *stab2* guide 3, alt | sgRNA template | taatacgactcactataGGATACGCTAAGATGCCACCgttttagagctagaa | This paper |
| oGMB541 | *stab2* guide 4, alt | sgRNA template | taatacgactcactataGGATTCGAGAAGTTTGTTGAgttttagagctagaa | This paper |
| oGMB116 | *slc24a5* guide 1 | sgRNA template | taatacgactcactataGGCACCGTGAAGAATCCTTCgttttagagctagaa | This paper |
| oGMB117 | *slc24a5* guide 2 | sgRNA template | taatacgactcactataGGTGACGGCGGCGACACTGAgttttagagctagaa | This paper |
| oGMB119 | *slc24a5* guide 3 | sgRNA template | taatacgactcactataGGTTGACGTGGGCTCCCCGGgttttagagctagaa | This paper |
| oGMB120 | *slc24a5* guide 4 | sgRNA template | taatacgactcactataGGGTGTCTCCTGCTGGTGTAgttttagagctagaa | This paper |
| KE stab2 fw primer | Genotype *stab2(ibl2)*, forward | Genotyping | GCATCAAAACAAATGTAACACAGC | This paper |
| KE stab2 rv primer | Genotype *stab2(ibl2)*, reverse | Genotyping | ATACACACAGCGGGTAGAGC | This paper |
| oGMB572 | Non-barcoded forward (oGMB279 site) | NGS sequencing | atacataaaagtgatgcccctg | This paper |
| oGMB573 | Barcode 1 reverse (oGMB279 site) | NGS sequencing | TCAGCTAAGGTAAATGGGCTTTATGATGTCACACC | This paper |
| oGMB574 | Barcode 2 reverse (oGMB279 site) | NGS sequencing | TCAGTGAGCGGAAATGGGCTTTATGATGTCACACC | This paper |
| oGMB575 | Barcode 3 reverse (oGMB279 site) | NGS sequencing | TCAGTCTATTCGTATGGGCTTTATGATGTCACACC | This paper |
| oGMB576 | Non-barcoded forward (oGMB280 site) | NGS sequencing | ctggaattcagcatggaaaatgttc | This paper |
| oGMB577 | Barcode 1 reverse (oGMB280 site) | NGS sequencing | TCAGCTAAGGTAActaatacCTTGATCACAATGAATTCC | This paper |
| oGMB578 | Barcode 2 reverse (oGMB280 site) | NGS sequencing | TCAGTGAGCGGAActaatacCTTGATCACAATGAATTCC | This paper |
| oGMB579 | Barcode 3 reverse (oGMB280 site) | NGS sequencing | TCAGTCTATTCGTctaatacCTTGATCACAATGAATTCC | This paper |
| oGMB580 | Non-barcoded forward (oGMB281 site) | NGS sequencing | GTGTGAACCTGTCAATCAGTG | This paper |
| oGMB581 | Barcode 1 reverse (oGMB281 site) | NGS sequencing | TCAGCTAAGGTAAgacataaattaagatgtattacCTGCAGT | This paper |
| oGMB582 | Barcode 2 reverse (oGMB281 site) | NGS sequencing | TCAGTGAGCGGAAgacataaattaagatgtattacCTGCAGT | This paper |
| oGMB583 | Barcode 3 reverse (oGMB281 site) | NGS sequencing | TCAGTCTATTCGTgacataaattaagatgtattacCTGCAGT | This paper |
| oGMB584 | Non-barcoded forward (oGMB282 site) | NGS sequencing | CGGCACATATTGTGAAGgtattc | This paper |
| oGMB585 | Barcode 1 reverse (oGMB282 site) | NGS sequencing | TCAGCTAAGGTAAaaactaactgtagatgtagacattc | This paper |
| oGMB586 | Barcode 2 reverse (oGMB282 site) | NGS sequencing | TCAGTGAGCGGAAaaactaactgtagatgtagacattc | This paper |
| oGMB587 | Barcode 3 reverse (oGMB282 site) | NGS sequencing | TCAGTCTATTCGTaaactaactgtagatgtagacattc | This paper |
| oGMB293 | Generate runx1+23 lifeact-mCherry, forward | Gibson cloning | tgctttcgaatttcttgatcaaatctgcgacacccatgggatgtctgtttctgaggttgc | This paper |

| | | | | |
|---|---|---|---|---|
| oGMB294 | Generate runx1+23 lifeact-mCherry, reverse | Gibson cloning | gatcaagaaattcgaaagcatctcaaaggaagaagtgagcaagggcgaggag | This paper |
| | *stab2* reverse† | WISH probe synthesis | AAAGAGAGCTGCACCGACT | Hagedorn *et al, Developmental Cell,* 2023 |
| | *stab2* forward* | WISH probe synthesis | TTGTGGATTACGGGGTTCGG | Hagedorn *et al, Developmental Cell,* 2023 |
| | *stab1* reverse† | WISH probe synthesis | CGCCGTTCTATAATGCACCG | Hagedorn *et al, Developmental Cell,* 2023 |
| | *stab1* forward* | WISH probe synthesis | AAGGCGTACTATGTCCTCAGGC | Hagedorn *et al, Developmental Cell,* 2023 |
| | *mrc1a* reverse† | WISH probe synthesis | ACGGCATTCCACAAACCAGA | Hagedorn *et al, Developmental Cell,* 2023 |
| | *mrc1a* forward* | WISH probe synthesis | GTGTCCCCTCATCAATGCCA | Hagedorn *et al, Developmental Cell,* 2023 |
| oGMB244 | *dab2* reverse† | WISH probe synthesis | gcacaggaatacttgcatagaggaa | This paper |
| oGMB243 | *dab2* forward* | WISH probe synthesis | cctcggggagaaggagagta | This paper |
| oGMB406 | *sele* reverse† | WISH probe synthesis | CATGGCTGCACATCACTGTC | This paper |
| oGMB405 | *sele* forward* | WISH probe synthesis | AAGCAAGGGATCTCACCACA | This paper |

*The T3 sequence CATTAACCCTCATAAAGGGAA was added to the 5' end of each forward primer for WISH probe synthesis

†The T7 sequence TAATACGACTCACTATAGGG was added to the 5' end of each reverse primer for WISH probe synthesis.

**Table S2 | Plasmids**

| Plasmid name | Description | Source | Notes |
|---|---|---|---|
| pGMB56 | lifeact-mCherry driven by runx1+23 enhancer and minimal promoter | This paper | Used to generate *Tg(runx1:lifeact-mCherry)* |
| pZSI3 | hLNGFR-p2A-mCherry driven by runx1+23 enhancer and minimal promoter | This paper | Construct from which pGMB56 was generated using Gibson cloning |
| GW runx1+23-hLNGFR-mly7GFP | hLNGFR driven by runx1+23 enhancer and minimal promoter | Wattrus & Zon, Zebrafish, 2020[85] | Construct from which pZSI3 was generated using Gibson cloning |

**Table S3 | RNAseq FPKM values from selected hematopoietic genes**

| Name | Gene ID | *runx(-)cd41(+)* | *runx(+)cd41(+)* | *runx(+)cd41(-)* | Chromosome | Gene start | Gene end | Strand |
|---|---|---|---|---|---|---|---|---|
| *runx1* | ENSDARG00000087646 | 1.78093232 | 6.12817683 | 0.5393557 | 1 | 1362932 | 1402303 | - |
| *itga2b* | ENSDARG00000018687 | 0 | 27.8554782 | 0.15943671 | 3 | 21350509 | 21382491 | - |
| *myb* | ENSDARG00000053666 | 4.56910884 | 30.9095758 | 0.79415534 | 23 | 31815423 | 31830043 | + |
| *gata2b* | ENSDARG00000009094 | 4.44914243 | 15.1279357 | 0.85745142 | 6 | 40794015 | 40803800 | + |
| *ptprc* | ENSDARG00000071437 | 6.48797882 | 14.5598439 | 1.7320793 | 22 | 22977268 | 23067859 | - |
| *blf* | ENSDARG00000043126 | 21.9869739 | 356.173419 | 55.6438925 | 22 | 9917007 | 9923692 | + |
| *cebpa* | ENSDARG00000036074 | 69.6925813 | 106.935946 | 49.3083478 | 7 | 38084986 | 38087865 | - |
| *gata1a* | ENSDARG00000013477 | 0.12575285 | 9.78311668 | 31.2481491 | 11 | 25411914 | 25418856 | - |
| *hbbe2* | ENSDARG00000045143 | 0 | 6.85073268 | 1398.89306 | 12 | 20349865 | 20350629 | - |
| *hbbe1.1* | ENSDARG00000113599 | 70.7113449 | 2229.80796 | 100811.873 | 3 | 55114097 | 55114874 | + |
| *hbae3* | ENSDARG00000079305 | 46.7206209 | 1351.7368 | 97094.508 | 3 | 55146418 | 55147788 | - |
| *cahz* | ENSDARG00000011166 | 4.18922798 | 85.8433489 | 6908.42985 | 2 | 29477383 | 29485408 | - |
| *spi1b* | ENSDARG00000000767 | 56.4962101 | 75.6804271 | 16.3035788 | 7 | 32644673 | 32659353 | - |
| *mpeg1.1* | ENSDARG00000055290 | 2.15145279 | 8.49601485 | 3.72757449 | 8 | 29710110 | 29713621 | - |
| *mfap4* | ENSDARG00000090783 | 16.9606702 | 66.2654009 | 20.0949865 | 1 | 59090583 | 59092056 | + |
| *irf8* | ENSDARG00000056407 | 3.20708921 | 11.9951888 | 2.71051423 | 18 | 30567945 | 30573849 | + |
| *mpx* | ENSDARG00000019521 | 256.767364 | 204.546763 | 41.0180443 | 10 | 7778870 | 7785945 | - |
| *lyz* | ENSDARG00000057789 | 5.68244408 | 17.9837621 | 2.18425643 | 24 | 24035863 | 24038800 | - |
| *cpa5* | ENSDARG00000021339 | 249.064635 | 288.173385 | 51.9479668 | 25 | 18459924 | 18470695 | - |
| *mpl* | ENSDARG00000039222 | 0.11192271 | 3.55822323 | 0.10168744 | 6 | 33949280 | 33980881 | - |
| *rag1* | ENSDARG00000052122 | 0.41290322 | 1.32631978 | 0.30894167 | 25 | 9003230 | 9009902 | + |
| *rag2* | ENSDARG00000052121 | 0.44224494 | 1.91139192 | 0.4018019 | 25 | 9010425 | 9013963 | - |
| *ikzf1* | ENSDARG00000013539 | 14.725027 | 42.0403549 | 19.5771054 | 13 | 15957508 | 15994472 | - |
